# Global organization of phenylpropanoid and anthocyanin pathways revealed by proximity labeling of trans-cinnamic acid 4-hydroxylase (CYP73A412) in *Petunia inflata* petal protoplasts

**DOI:** 10.1101/2024.04.05.588085

**Authors:** Javiera Aravena-Calvo, Silas Busck-Mellor, Tomas Laursen

## Abstract

The phenylpropanoid pathway is one of the main carbon sinks in plants, channeling phenylalanine towards thousands of products including monolignols, stilbenes, flavonoids and volatile compounds. The enzymatic steps involved in many of these pathways are well characterized, however the physical organization of these enzymes within the plant cell remains unknown Proximity-dependent labeling allows untargeted determination of protein interactions *in vivo*, and therefore stands as an attractive alternative to targeted binary assays for determining global protein-protein interaction networks. In this study, we show a TurboID-based proximity labeling system developed to study protein interaction networks of the core phenylpropanoid pathway in petunia. Here, the endoplasmic reticulum (ER) membrane anchored cytochrome P450 cinnamic acid 4-hydroxylase (C4H, CYP73A412) from *Petunia inflata* was coupled to TurboID and expressed in protoplasts derived from anthocyanin-rich petunia petals. Potential interactors were isolated using streptavidin beads, digested and quantified by mass spectrometry. Among the enriched proteins, we identified multiple soluble enzymes from the late anthocyanin pathway, other CYP73 isoforms, as well as additional ER membrane anchored CYPs including *p*-coumaric acid 3-hydroxylase (C3’H, CYP98A2). Our results suggest that CYP73A412 co-localizes with enzymes from the phenylpropanoid- and downstream anthocyanin pathways, supporting the idea that CYP73s may serve as ER anchoring points for these metabolic pathways. Moreover, this study demonstrates the feasibility of using protoplasts to perform global mapping of protein network for enzymes in their native cellular environment.

## INTRODUCTION

Phenylpropanoids are specialized metabolites derived from L-phenylalanine, present in land plants. The phenylpropanoid pathway branches out to produce several major compound classes such as monolignols, stilbenes, coumarins, volatile phenylpropanoids and flavonoids, including flavonols and anthocyanins. Anthocyanins are pigments that accumulate in different plant tissues and act as photoprotectants by absorbing UV light and scavenging free radicals (Guo et al., 2008). Additionally, anthocyanin pigments provide a palette of colors ranging in color from orange to purple, pink and blue that serve to attract pollinators and seed dispersers to specialized plant tissues such as flowers and fruits (Winkel-Shirley, 2001). This has made the phenylpropanoid pathway one of the best characterized pathways in plant specialized metabolism. The ability to guide phenylalanine towards specific downstream products on-demand, combined with experimental data from binary protein-protein interaction assays has prompted the hypothesis that the pathway may organize into dynamic enzyme complexes termed metabolons (Knudsen et al., 2018; Laursen et al., 2015; Nakayama et al., 2019; Winkel, 2004). Furthermore, evidence from cyanogenic glucoside biosynthesis suggests that enzymes of the cytochrome P450 superfamily, which are anchored to the endoplasmic reticulum (ER) membrane, serve as assembly points for metabolons (Jørgensen et al., 2005; Laursen et al., 2021).

The general phenylpropanoid pathway (Figure 1) converts phenylalanine to 4-coumaroyl-CoA by the action of a soluble phenylalanine ammonia-lyase (PAL), an ER membrane anchored P450 enzyme, cinnamic acid 4-hydroxylase (C4H, CYP73A412), and a soluble 4-coumaryl-CoA ligase (4CL). This pathway remains one of the few that has been proven to form a metabolon that involve protein-protein interactions and substrate channeling (Achnine et al., 2004; Rasmussen & Dixon, 1999). Substrate channeling has been demonstrated by isotopic dilution experiments in tobacco cell cultures and microsomes. Here, the high proportion of [^3^H]-4-coumaric acid converted from the fed [^3^H]-L-phenylalanine did not equilibrate with the [^14^C]-4-coumaric acid originated from feeding the intermediate [^14^C]-*trans*-cinnamic acid, showing the rapid channeling of [^2^H]-*trans*-cinnamic acid from PAL to CYP73A412 (Rasmussen & Dixon, 1999). Subsequent work using fluorescence resonance energy transfer coupled with fluorescence lifetime imaging microscopy (FRET-FLIM) showed that tobacco CYP73A412 interacts stronger with PAL1 compared to PAL2 (Achnine et al., 2004). Taken together, these findings support the formation of a metabolon in the early phenylpropanoid pathway.

**Figure 1.**
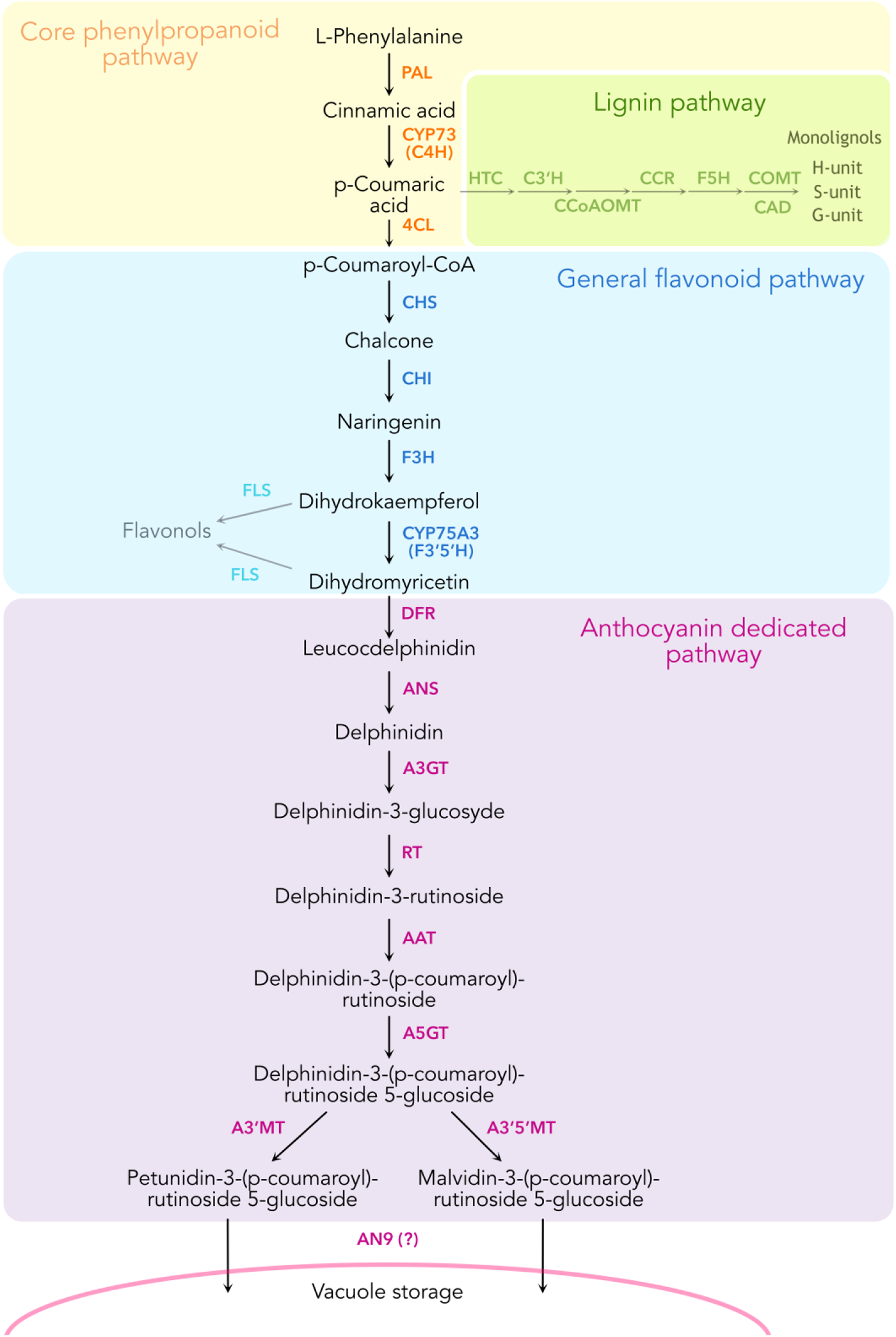
Schematic representation of the enzymatic reactions leading to the biosynthesis of anthocyanins in *Petunia inflata*, showing some of the branching pathways such as lignin and flavonol biosynthesis.

To date only a handful of metabolons have been experimentally proven (Laursen et al., 2016; Mucha et al., 2019; Zhang et al., 2018). This is largely due to the lack of proper techniques for elucidation of global protein interaction networks *in planta*. The recent development of proximity-dependent labeling has proven a powerful tool for the identification of protein networks in an untargeted manner that is gaining popularity also in the plant sciences (Xu et al., 2023). Biotin-dependent proximity labeling is conducted by fusing a highly active biotin ligase to a protein of interest (bait), which is expressed either stably or transiently in the organism. The addition of the biotin substrate allows the ligase to activate biotin and enable covalent coupling to lysine residues of all proteins in proximity to the bait. Biotinylated proteins are then captured by affinity purification, digested into peptides and identified by mass spectrometry (MS). One strength of proximity labeling is its ability to detect weak and transient interactions, which are frequently lost by other methods such as co-immunoprecipitation or tandem affinity purification. Furthermore, proximity labeling may provide spatiotemporal resolution of the protein interaction networks of a given biological process under native conditions by controlling labeling time (Branon et al., 2018; Gingras et al., 2019; Yang et al., 2021). Among the recently developed biotin ligases, TurboID is the most efficient of the biotin ligases at room temperature and therefore displays faster labeling times, making it appropriate for studying dynamic interactions in plants (Arora et al., 2020; Mair et al., 2019; Zhang et al., 2019). TurboID has revealed protein interaction networks of regulators involved in NLR immune response in tobacco (Zhang et al., 2019) and interactomes of endocytic TPLATE protein complex in *Arabidopsis* and tomato (Arora et al., 2020). Moreover, it has allowed the elucidation of proteomes from cell-type specific compartments in *Arabidopsis* guard cells without the challenge of isolating them (Mair et al., 2019).

In this study, we develop transient TurboID-based proximity labeling to elucidate protein networks in the anthocyanin pathway. Using protoplasts from anthocyanin-rich flower petals from *Petunia inflata*, which provide a fast and reliable transformation system, we demonstrate multiplexed proximity labelling experiments without the need to generate stable mutant plant lines. By using this system with CYP73A412 as bait we found enrichment of several cytosolic enzymes from the late anthocyanin pathway. Our results suggest that these soluble enzymes localize on the ER surface near CYP73A412, indicating the presence of higher order metabolic complexes in the anthocyanin pathway, which may be anchored by interactions with CYP73A412. This is the first study reporting P450 protein networks elucidated by proximity labeling and stands as the basis for future experiments investigating metabolon formation in anthocyanin biosynthesis.

## RESULTS

### Global proteome profile of *Petunia inflata* petal protoplasts

Proximity labeling has been established in a few plant species, either by generating stable transgenic plants overexpressing bait-TurboID fusion constructs or by transient agroinfiltration in leaves (Arora et al., 2020; Mair et al., 2019; Zhang et al., 2019). Although agroinfiltration is possible directly in petunia petals (Verweij et al., 2008), the tissue does not tolerate the successive rounds of infiltration, which are required to introduce biotin post transfection. To overcome this, we sought to develop a workflow for TurboID-based proximity labeling using protoplasts isolated from petal tissues.

Petunia petals have a laminar morphology, which consists of an outer layer of pigmented epidermal cells sandwiching a few mesophyll cell layers (Cavallini-Speisser et al., 2021). To confirm that the enzymes of the general phenylpropanoid and flavonoid pathways were present in protoplasts isolated from mature flower petals, we performed global proteomics on protoplasts isolated from mid-development flower buds (stage 3, “S3”) and newly open flowers (stage 5, “S5”). After filtering and imputation, we obtained 11135 proteins quantified in both stages. Enzymes involved in the biosynthesis of anthocyanins were slightly more abundant in protoplasts from S3 petals compared to protoplasts from S5 petals, as seen in the ranked protein abundances for both stages (Figures 2A and 2B). Nevertheless, analysis of differential enrichment of proteins between S5 and S3 showed that abundances of most of the biosynthetic enzymes of the anthocyanin pathway were not significantly different in the two stages, with the exception of CHSa and a CYP73A412 paralog, C4Hc, which were significantly enriched in S3 (Figure 2C). Because anthocyanin biosynthetic enzymes are still among the most abundant proteins in S5 protoplasts, and flowers from this stage provided a much greater yield of protoplasts than flowers from S3, we used S5 protoplasts for further experiments.

**Figure 2:**
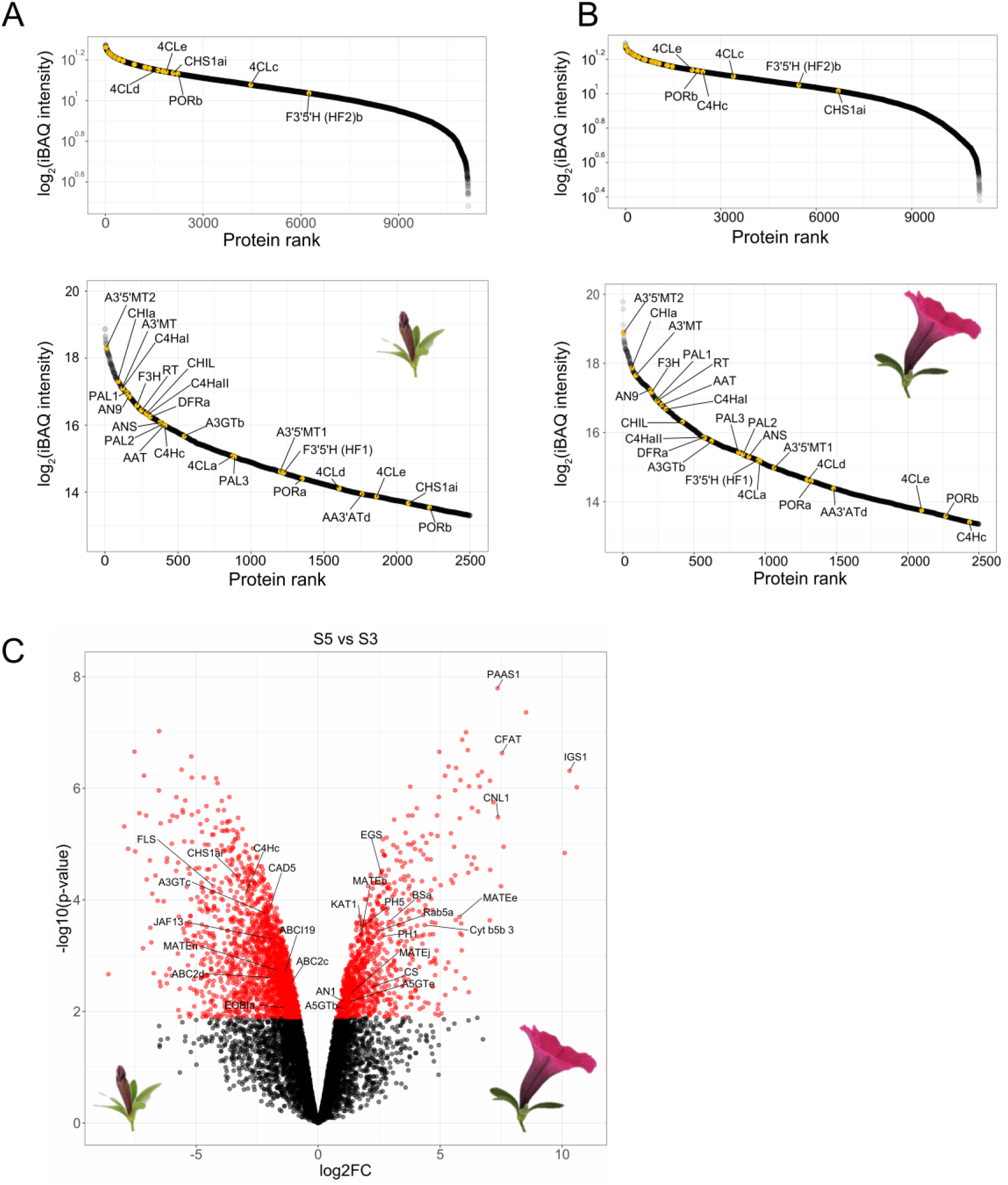
Global proteomics of protoplasts derived from *P. inflata* petals. **(A)** Rank abundance curve of total expressed proteins in protoplasts from petals at stage 3, upper panel corresponds to the ranking of the total number of proteins, whereas lower panel shows the 2500 most abundant proteins in protoplasts from S3 flowers. **(B)** Rank abundance curve of total expressed proteins in protoplasts from petals at stage 5, upper panel corresponds to the ranking of the total number of proteins, whereas lower panel shows the 2500 most abundant proteins in protoplasts from S5 flowers. **(C)** Volcano plot of differentially enriched proteins in S5 vs S3. Red circles to the left correspond to significantly enriched proteins in S3, red circles in the right are proteins significantly enriched in S5. Log2FC, Log2 fold-change.

### Development of proximity labeling in petunia protoplasts

We developed a methodology for performing TurboID-based proximity labeling in *P. inflata* petal protoplasts, shown in Figure 3A. Due to the highly conserved N-terminal transmembrane domain of eukaryotic P450s, TurboID was fused to the C-terminal of CYP73A412 including a short flexible linker (SGGGGGSGGG) between the proteins (Figure 3B). Petal protoplasts were transformed with expression vectors encoding the control construct p35S:ER-TurboID-EGFP, consisting of TurboID fused N-terminally to the transmembrane anchor domain from the cytochrome P450 CYP51 and C-terminally to EGFP, and the bait construct p35S:CYP73A412-TurboID (Figure 3C).Transformation efficiency of protoplasts ranged between 20-40% in all our experiments. To confirm correct ER localization and verify expression in transformed protoplasts we expressed a construct containing CYP73A412-TurboID fused to EGFP (p35S:CYP73A412-TurboID-EGFP). Untransformed protoplasts (WT) were included as control for endogenously biotinylated proteins. After overnight incubation, we observed fluorescence in protoplasts transformed with both ER-TurboID-EGFP and CYP73A412-TurboID-EGFP fusion proteins, with fluorescence limited to membrane network structures surrounding the nucleus, characteristic for ER localized proteins (Figure 4A). These results indicate that the fusion proteins were successfully expressed in this protoplast system and that the expressed proteins were correctly localized to the ER membrane.

**Figure 3.**
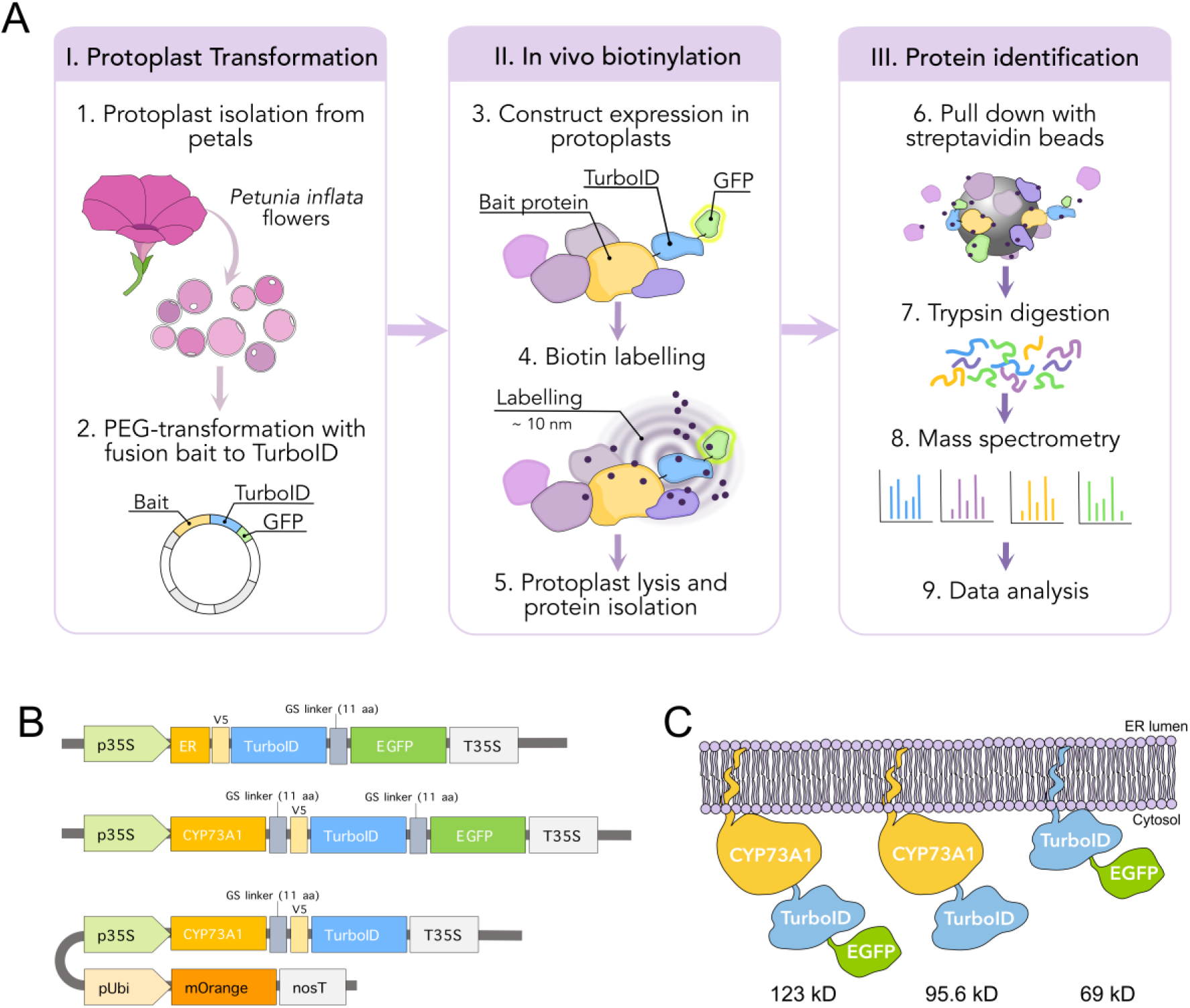
Design and workflow of proximity labeling in protoplast from P. inflata. **A)** Vector design for control ER-TurboID-EGFP and baits CYP73A1-TurboID-EGFP and CYP73A1-TurboID. The CYP73A1-TurboID construct contains a cassette for expression of mOrange as a reporter for transformation. **B)** Diagrammatic representation of the fusion proteins and their molecular weights. **C)** Workflow of the proximity labelling procedure. I, protoplasts are isolated from Petunia freshly opened flowers and PEG-transformed with plasmids containing TurboID control or bait-TurboID fusion. II, After assessment of transformation efficiency, biotinylation is done *in vivo* by incubating the protoplasts with 50 µM biotin solution for 0 and 3 hours, after which the protoplasts are washed three times. III, Protoplasts are lysed, proteins extracted and affinity purified for subsequent identification by mass spectrometry.

**Figure 4.**
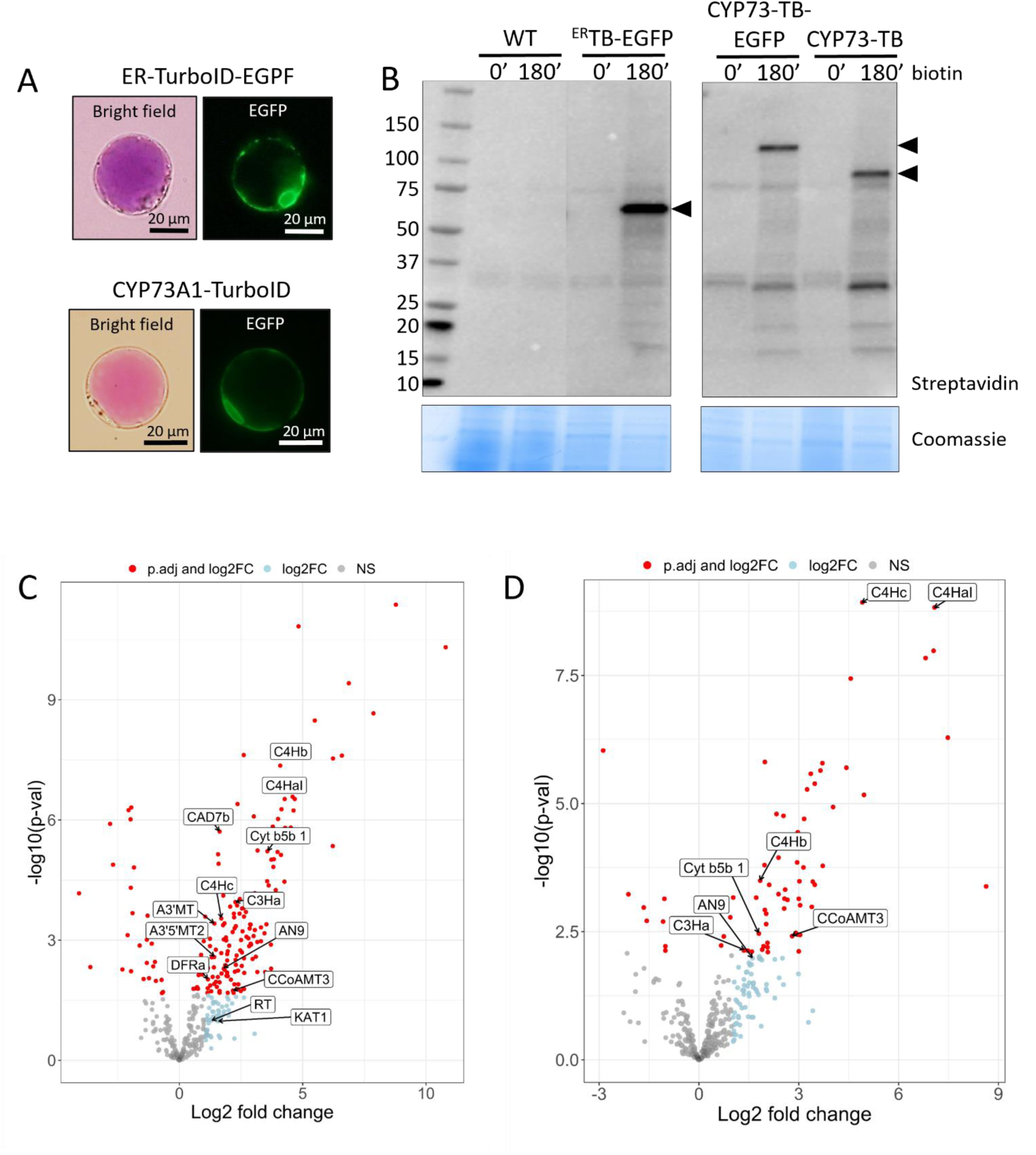
Establishment of proximity labeling in *P. inflata* petal protoplasts. **A)** Subcellular localization of ER-TurboID-EGFP (upper panel) and CYP73A1-TurboID-EGFP in protoplasts isolated from petal epidermis (lower panel). **B)** Expression and activity of ER-TurboID-EGFP, CYP73A1-TurboID-EGFP and CYP73A1-TurboID were analyzed by immunoblots using streptavidin-HRP antibody (upper panel). Coomassie blue staining is showed as loading control. Points of arrow indicate the strongest bands corresponding to cis-biotinylation, ER-TurboID-EGFP: 69 kD; CYP73A1-TurboID-EGFP: 123 kD; CYP73A1-TurboID: 95.6 kD. **C)**, **D)** Volcano plots showing log2-fold change and –log10(p-value) values from differential expression analysis comparing CYP73A1-TurboID vs WT at different biotin incubation timepoints: t=0 minutes (**C**) and t=180 minutes (**D**). Proteins significantly enriched (adjusted p-value < 0.05) in CYP73A1-TurboID are showed as red circles on the right half of each plot. Textboxes indicate the names of enzymes involved in the biosynthesis of anthocyanins: A3’MT: Anthocyanin 3’-methyltransferase; A3’5’MT2: Anthocyanin 3’5’-methyltransferase; AN9: glutathione transferase; C4HaI, C4Hb, C4Hc: CYP73A paralogs; C3Ha: cinnamate 3-hydroxylase a; CAD7: cinnamyl alcohol dehydrogenase 7a; CCoAMT3: Caffeoyl-CoA O-methyltransferase 3; Cyt b5b1: Cytochrome b5b 1; DFR: dihydroflavonol reductase; KAT1: 3-ketoacyl-CoA thiolase 1; RT: anthocyanin rhamnosyltransferase.

To investigate the ability of TurboID fusions to biotinylate proteins in our protoplast system, we incubated transformed protoplasts with 50 µM biotin for 0 and 3 h. Next, we extracted proteins and analyzed the pattern of biotinylation by streptavidin immunoblot to evaluate the expression of catalytical active TurboID. Incubation with biotin for 3 h induced strong self-labeling (cis-biotinylation) in protoplasts expressing ER-TurboID-EGFP (69 kD), CYP73A412-TurboID-EGFP (123 kD) and CYP73A412-TurboID (95.6 kD) whereas only endogenously biotinylated proteins were observed in the absence of biotin (t =0) and untransformed controls (WT) (Figure 4B). Besides cis-biotinylation, trans-biotinylation was evident in protoplasts expressing TurboID-fusions visualized by a characteristic smear on the immunoblots (Arora et al., 2020; Mair et al., 2019), which corresponds to proteins labeled by encountering TurboID fusions inside the cell.

To eliminate interference of EGFP on CYP73A412 interactions, subsequent proximity labeling experiments was performed using the bait CYP73A412-TurboID without EGFP.

### Identification of proteins neighboring CYP73A412

Biosynthesis of phenylpropanoids is expected to occur at the cytosolic surface of the ER membrane, and CYP73A412 has been proposed to act as a hotspot to coordinate assembly of soluble enzymes for production of phenylpropanoid derivatives (Winkel-Shirley, 1999). To investigate interaction networks around CYP73A412, we purified proteins that were biotinylated after expressing CYP73A412-TurboID. The amount of biotinylated proteins obtained was highly variable among the different treatments (Supplementary Figure 1A), however these quantities were enough to quantify peptides by mass spectrometry. Comparison of proteins enriched in CYP73A412-TurboID protoplasts versus untransformed and TurboID-EGFP controls at 0 and 3 h after biotin incubation by label-free quantification (LFQ) identified 424 proteins after removing proteins with too many missing values. Principal component analysis (PCA) (Supplementary Figure 1B) showed a clear separation between CYP73A412-TurboID and control groups. ER-TurboID-EGFP grouped with WT controls at both timepoints, suggesting that the protein compositions and quantities of these groups were very similar. The data set from CYP73A412-TurboID was first contrasted against WT to identify enriched proteins (possible interactors) and background proteins consisting ofnaturally biotinylated proteins such as Acetyl-CoA carboxylases (ACCases), and biotin carboxyl carrier protein (BCCP) subunits, as well as proteins binding non-specifically to the beads. Pair-wise comparison of CYP73A412-TurboID and ER-TurboID-EGFP did not result in enriched proteins, consistent with the clustering of WT and ER-TurboID-EGFP seen in the PCA (Supplementary Figures 1, 3).

After pair-wise comparison of the bait with WT samples, we ended up with 185 significantly enriched proteins in protoplast samples before biotin incubation (t= 0) and 69 proteins in the biotin treatment (t=3h) (Figure 4C and 4D). Most of the proteins identified were annotated for processes associated to mitochondrial metabolism, translation machinery and chaperones (Supplementary Tables 2 and 3). However, among significantly enriched proteins we found additional two CYP73A412 isoforms (C4Hb (CYP73A413), C4Hc (CYP73A414) and soluble enzymes from different branches of the phenylpropanoid pathway including the monolignol branch and anthocyanin branch. From the anthocyanin pathway, we identified dihydroflavonol synthase (DFR), anthocyanin rhamnosyltransferase (RT), the anthocyanidin 3’ and anthocyanidin 3’5’ O-methyltransferases (A3’MT and A3’5’MT) that perform the final methylathion of delphinidin to form the predominant anthocyanins petunidin and malvidin (Provenzano et al., 2014), and a glutathione transferase (AN9), which has been suggested to be involved in the transport of anthocyanins to the vacuole (Alfenito et al., 1998; Buhrman et al., 2022; Mueller et al., 2000), but was recently shown to participate in the conversion of leucoanthocyanidins to anthocyanidins together with anthocyanidin synthase (Eichenberger et al., 2023). From the monolignol pathway, we found the soluble enzymes cinnamyl alcohol dehydrogenase (CAD), caffeoyl-CoA O-methyltransferase (CCoAOMT), and the membrane anchored *p*-coumarate 3-hydroxylase (C3’H, CYP98A2).

## DISCUSSION

### *P. inflata* petal protoplasts provide a versatile system for proximity labeling experiments

Proximity labeling has proven to be a powerful tool to elucidate subcellular proteomes and protein-protein interaction networks. It stands out from traditional binary protein-protein interaction methods, such as yeast two-hybrid, bi-molecular fluorescence complementation (BiFC) and FRET methods, in providing an untargeted approach for studying protein-protein interactions in native conditions and can be used to detect weak or transient interactions that may be lost by other untargeted methods such as co-immunoprecipitation (Branon et al., 2018). To date, only a handful of studies have successfully reported biotin ligase-based proximity labeling in model plant species such as tobacco, *Arabidopsis* and tomato (Arora et al., 2020; Feng et al., 2023; Gryffroy et al., 2023; Huang et al., 2020; Khan et al., 2018; Mair et al., 2019; Zhang et al., 2019), mostly involving stable transformation of plants to expressing the bait of interest. Here, we demonstrate proximity labeling in *P. inflata,* which is an important model for anthocyanin biosynthesis and one of the few petunias with a sequenced genome (Bombarely et al., 2016). Here, we provide a protocol for studying protein interactions in the biosynthesis of anthocyanins by transient expression in protoplasts from highly specialized cell types as the flower petals. Protoplasts have been extensively used to study biological processes such as light responses, signal transduction, protein interactions, and multi-omics (Sheen, 2001; Xu et al., 2022). Faraco et al. (2011) developed a protocol for isolation and transformation of petunia petal protoplasts with high transformation efficiency (up to 60%), which enabled the study of subcellular processes related to anthocyanin accumulation and vacuole acidification (Faraco et al., 2017; Li et al., 2021). Here, we show that transient expression in petunia protoplasts makes a versatile alternative to stable transformation for performing proximity labeling, with fast experimental turnaround.

The temporal expression of biosynthetic genes from the anthocyanin pathway during flower development has revealed that anthocyanin biosynthesis peaks at stage 3 and decrease towards the end of the bud development and flower opening (Quattrocchio et al., 1993, 1999). Therefore, we were concerned that the anthocyanin pathway would already be downregulated in S5 flowers, which was used for preparation of protoplasts followed by proximity labeling. However, while our global proteomics analysis showed that enzymes belonging to the phenylpropanoid and anthocyanin pathways are among the most abundant proteins in protoplasts from stage S3 (Figure 2A), the biosynthetic enzymes remained highly abundant in protoplasts from open flowers (S5) (Figure 2B), similar as reported by Prinsi et al. (2016). These findings contrast with previous studies based on transcript levels (Quattrocchio et al., 1993, 1999). Such discrepancies between gene expression measured at the transcript versus protein levels have previously been reported in plant systems (Mergner & Kuster, 2022), indicating that direct measurement of protein levels by mass spectrometry is preferable to determine the optimal tissues to perform proximity labeling experiments.

### Considerations for proximity labeling experiments in *P. inflata* petal protoplasts

Plant protoplasts are cells where the cell wall has been removed and therefore can be comparable to mammalian cells due to the similar exposed cellular membrane (Sheen, 2001). Consequently, we chose concentrations of biotin (50 µM) typically used for proximity labeling experiments in HEK 293T cells (Branon et al., 2018) and in rice protoplasts (Lin et al., 2017). Our data suggests that endogenous biotin levels (t=0) were sufficient for TurboID-mediated biotinylation to provide reliable protein quantification (Figures 4A and 4B), even though biotinylation was undetectable in our immunoblots. We speculate that relying solely on labeling with endogenous biotin may reduce background and lead to more accurate identification of interactors, though this effect remains to be systematically assessed. Two of the most challenging aspects of working with protoplasts relates to yields and transformation efficiency. Hence, approaches that obviate the requirement for biotin treatments would significantly decrease experimental complexity by reducing sample number and enable higher throughput in large-scale proximity labeling experiments. The main drawback of this approach is that long labeling times required might reduce temporal resolution, and so may not be suitable for experiments to determine temporally dependent interactions.

### A glimpse into the CYP73A412 interactome

Protein-protein interactions among enzymes in the anthocyanin pathway have been investigated in several plant species such as rice, *Arabidopsis*, soybean, snapdragon and torenia (Fujino et al., 2018; Nakayama et al., 2019), and these studies clearly demonstrate that biosynthesis of flavonoids involves formation of different enzyme complexes. Nevertheless, the exact functional consequences of such protein complexes are not fully understood, and their exact compositions do not seem to be conserved between different species (Nakayama et al., 2019).

CYP73A412 catalyzes the conversion of cinnamic acid to *p*-coumaric acid, an essential early step in the phenylpropanoid pathway (Vogt, 2010). Because *p-*coumaric acid is a key branch point metabolite serving as substrate for production of coumarins, mono lignols, stilbenoids, curcuminoids, floral volatiles and flavonoids, we speculate that CYP73A412 may control downstream production through intricate interaction networks with dedicated enzymes for individual metabolic branches. Using CYP73A412 as a bait has revealed several potential interactors from the routes branching out the general phenylpropanoid pathway, essentially downstream enzymes from the lignin and anthocyanin pathways, supporting the role of P450s as hub for metabolic complexes (Jørgensen et al., 2005; Laursen et al., 2015; Ralston & Yu, 2006; Zhang & Fernie, 2021). Though the interactions observed in this study remains to be verified by complementary approaches, our results indicate new associations between CYP73A412 and enzymes from both anthocyanin (DFR, A3’MT, A3’5’MT and AN9), and monolignol pathways (C3’H, CAD, CCoAMT) in petunia (Figures 4A and 4B).

Our results are consistent with previous studies revealing interactions between CYP73A412 and the initial enzyme of the lignin pathway, CYP98A3, in *Arabidopsis* (Bassard et al., 2012). More recently, a combination of FRET and yeast two hybrid (Y2H) experiments suggested that CYP73A5 (a C4H paralog), CYP98A3 (C3’H) and CYP84A1 (F5H) are in close vicinity and co-localize in the ER membrane of *Arabidopsis* cells, but do not interact directly (Gou et al., 2018). To our knowledge, this report is the first to suggest interactions between CYP73A412 and enzymes of the anthocyanin pathway. While the interactions remain to be validated by complementary methods, it is possible that the absence of such interactions in previous reports relate to the specific tissues studied or the sensitivity of the techniques to capture highly dynamic and low affinity interactions (Bassard et al., 2012; Gou et al., 2018).

Surprisingly, none of the previously known interactors of CYP73A412 were identified among the enriched proteins in our proximity labeling experiment, such as phenylalanine ammonia lyase (PAL), P450 oxidoreductase (POR), membrane steroid binding protein (MSBP) and 4-coumarate-CoA ligase (4CL) (Achnine et al., 2004; Bassard et al., 2012; Gou et al., 2018). Future combinatorial studies including more anthocyanin enzymes as bait will provide a more holistic view of the organization of the pathway.

## CONCLUSION

We demonstrate the use of proximity labeling in *P. inflata* protoplasts isolated from petals of S5 flowers. Using CYP73A412 as bait fused to TurboID, we identified enzymes from different branches of the phenylpropanoid pathway, supporting a central role of this enzyme as anchoring point for channeling of phenylalanine towards production of diverse compounds according to the specific needs of the plant cell.

This study demonstrates that protoplasts can be used to map protein networks in their native cellular environment using TurboID-based proximity labeling. This provides a new method to use proximity labeling to study protein interaction networks in specialized cell types, and may provide a blueprint for its use in plants that are otherwise recalcitrant to *Agrobacterium-* based transformation.

## MATERIAL AND METHODS

### Plant material and growing conditions

*Petunia inflata* seeds were obtained from the Amsterdam petunia germplasm collection (Strazzer et al., 2023) and germinated in pots under greenhouse conditions at 25 °C day and 22 °C night, with a photoperiod of 16h light/ 8h dark. For protoplast isolation, metabolomics and proteomics analysis, corolla tissue was collected from bud stage (S3) and freshly opened flowers (S5).

### Protoplasts isolation

Isolation of protoplasts was performed as described in Faraco et al., 2011. *P. inflata* flower buds at stage 3 (S3) and freshly opened flowers at stage 5 (S5) were collected (Suppl. Figure 1). The anthocyanin-rich epidermal layer was peeled from the petal and used for protoplast isolation. Briefly, the tissue was incubated overnight in TEX buffer (Gamborg B5 medium, 500 mg/L MES, 750 mg/L CaCl2, [2*H2O] 250 mg/L NH4NO3, and 0.4 M sucrose, pH 5.7), containing 0.2% Macerozyme R10 and 0.4 % Cellulase R10 (Duchefa). After 16 h, the protoplast suspension was filtered with a 100 µm cell strainer (Falcon) and washed three times with TEX buffer, with a 100 x g centrifugation step between washes. The protoplasts were counted using a hemocytometer and subsequently used for global proteomics and proximity labeling experiments.

### Total protein isolation from protoplasts

A volume of isolated protoplasts containing 5 x 10^5^ cells was taken in three replicates, pelleted by centrifugation at 100 x g and lysed with 150 µL RIPA buffer (50 mM Tris, 150 mM NaCl, 1 mM EDTA, 1% NP40, 0.1% SDS, 0.5% sodium deoxycholate) supplemented with protease inhibitor (cOmplete™, Roche), 1 mM DTT and 1 mM PMSF. The lysate was centrifuged at 15000 x g at 4°C to remove cell debris and the supernatant was transferred to a new protein Lo-Bind tube. One hundred µg of the total protein extract was used for mass spectrometry analysis.

### Sample preparation for global proteomics

Sample extracts were diluted with 1:1 with 2x SDS lysis buffer (10% SDS, 100 mM Tris pH 8.5), reduced with 5 mM tris(2-carboxyethyl)phosphine (TCEP) for 15 min at 55 °C, and alkylated with 20 mM chloroacetamide for 30 min at room temperature. Protein purification and digest was performed following the PAC protocol (Batth et al., 2019). Peptides were acidified with TFA (final concentration 1%) and purified using SDB-RPS StageTips (Kulak et al., 2014).

### LC-MS global proteomics

Peptides were separated on a 25 cm column Aurora Gen2, 1.7uM C18 stationary phase (IonOpticks) with either a Vanquish Neo HPLC system or an EASY-nLC 1200 HPLC (Thermo Scientific) coupled via a captive-spray source to a timsTOF pro2 (Bruker Daltonics) mass spectrometer operated in DIA-PASEF mode (Meier et al., 2020). Peptides were loaded in buffer A (0.1 % formic acid) and separated with a non-linear gradient of 2 – 35% buffer B (0.1

% formic acid, 99.9% acetonitrile) at a flow rate of 400 nL/min over 90 min. Total run time was 101min including washing phase. The column temperature was kept at 50° C during all LC gradients. MS acquisition was performed as described previously for R270.

### Global proteomics analysis

For global proteomics of petunia protoplasts, data-independent acquisition (DIA) was processed in library-free mode using DIA-NN software (Demichev et al., 2022), using the *Petunia inflata* proteome (version 1.0.1, from https://solgenomics.net/) containing all proteins translated from the available genome (Bombarely et al., 2016). Search settings were as follows: Protease trypsin, peptide length 7-35 residues, precursor m/z 300-1800, precursor charge 1-4, maximum missed cleavages 1, Met oxidation as variable modification (up to 2 allowed), carbamidomethylation of Cys as fixed modification and N-termin Met excision enabled. Match-between-runs (MBR) was enabled, and Robust LC was used as quantification strategy. The R package DIAgui (https://github.com/mgerault/DIAgui) was used to calculate LFQ and iBAQ values from the intensities reported by DIA-NN. The iBAQ values were filtered to keep only proteins quantified in all three replicates in at least one group and subsequently, missing values were imputed using quantile regression imputation of left-censored data (QRILC) method (Zhang et al., 2018), assuming missing values not at random. Differential analysis of protein expression was done in R using the DEP package (Zhang et al., 2018).

### Construction of plasmids for proximity labeling assays

Full-length CYP73A412 (Peinf101Scf00951g08008.1, C4HaI) was amplified from cDNA isolated from *P. inflata* corolla with the primers listed in Supplementary Table 1. The resulting PCR fragment was subsequently cloned in pJET and verified by sequencing. TurboID was amplified from Addgene plasmid #127350 (Mair et al., 2019), retaining the N-terminal V5 tag. For proximity labeling TurboID was fused at the C terminus of CYP73A412, including a 10 amino acid linker SGGGGGSGGG between the proteins. To generate CYP73A412-TurboID-EGFP and CYP73A412-TurboID fusions, CYP73A412, TurboID and EGFP were separately amplified by PCR using primers (Supp) that allowed 15-20 nucleotides overlap among the PCR fragments and the destination vector for a seamless fusion. To build the TurboID control for the ER compartment, the first 40 nucleotides corresponding to the ER transmembrane domain of CYP51 were amplified by PCR. The PCR fragments were fused and cloned into a binary vector (pPZP200 backbone), after the CaMV35S promoter (p35S) by Gibson Assembly using NEBuilder^®^ HiFi DNA Assembly Master Mix (NEB). The resulting plasmids containing p35S::CYP73A412-TurboID-EGFP, p35S::CYP73A412-TurboID and p35S::ER-TurboID-EGFP were subsequently verified by sequencing.

### Protoplast transformation

A total 1.4 x 10^6^ protoplast cells per sample were resuspended in MMM solution (0.5 M mannitol, 15 mM MgCl_2_, 0.1% MES). 20 µg of supercoiled plasmid was added to the protoplasts together with PEG solution (0.4 M mannitol, 0.1 M Ca(NO_3_)_2_, 40% polyethylene glycol (PEG 4000) and incubated at room temperature for 1 h. Then, the protoplasts were washed three times with TEX buffer, resuspended in 2 mL TEX after the last centrifugation and incubated in darkness at 26°C for 16 h.

### Microscopy/ subcellular localization

Transformed protoplasts expressing CYP73A412-TurboID-EGFP and ER-TurboID-EGFP were visualized by fluorescence microscopy (Nikon Eclipse Ni-U), 16h after transfection. GFP fluorescence was observed using excitation at 488 nm and emission at 514 nm.

### Proximity labeling experiments

Three biological replicates of each transformation with p35S::CYP73A412-TurboID, p35S::ER-TurboID-EGFP (subcellular compartment control) and with buffer (WT control, unstransformed) were incubated in 2 mL TEX buffer containing 50 µM biotin for 0 and 3 h at 25 °C. Protoplasts were washed 5 times with ice-cold W5 solution (154 mM NaCl, 125 mM CaCl_2_, 5 mM KCl, 5 mM glucose, 2 mM MES, pH adjusted to 5.7), 100 x g centrifugation at 4°C between washes and pelleted after the last wash. To obtain total protein extracts, the protoplast pellet was lysed with ice-cold 200 µL RIPA buffer supplemented with proteinase inhibitor, 1 mM DTT and 1 mM phenylmethylsulfonyl fluoride (PMSF). The solution was incubated on ice for 15 min and then centrifuged for 10 min, 4°C at 15000 x g. The supernatant was transferred to a protein Lo-Bind microcentrifuge tube (Eppendorf) and total protein was subjected to colorimetric quantification with Pierce™ BCA protein assay kit (Thermo Scientific).

### Western blot analysis

Total protein extract (10 µg) was separated by SDS-PAGE and transferred to PVDF membranes (BioRad). For detection of biotinylated proteins as result of TurboID activity, the membrane was blocked with 5% bovine serum albumin in PBS with 0.05% Tween-20 for 1 h and then incubated with HRP-conjugated streptavidin (1:20000, Thermo Fisher Scientific) with shaking for 1 h at room temperature or 4°C overnight. Blots were visualized using a GelDoc MP Imaging System (BioRad).

### Affinity purification of biotinylated proteins

To capture biotinylated proteins, 100 µL slurry of magnetic beads conjugated with streptavidin (Thermo Fisher) was equilibrated by washing with RIPA buffer 3 times and subsequently incubated overnight with 100 µg of total protein extracted from petunia protoplasts, under continuous rotation at 4°C. After incubation, the beads were washed as follows: 3x with cold RIPA buffer supplemented with 1 mM DTT for 2 min, 1x with cold 1 M KCl, 1x with cold 0.1 M Na_2_CO_3_, 1x with 2 M urea in 10 mM Tris-HCl pH 8, 2x with cold RIPA buffer supplemented with 1 mM DTT and 3x with cold PBS to remove traces of detergents. One tenth of the beads was boiled in Laemmli buffer containing 20 mM DTT and 2 mM biotin and used for immunoblots. The rest of the beads were kept dried at -80°C until further processing.

### On bead digest

Washed beads were incubated for 30 min with elution buffer 1 (2 M Urea, 50 mM Tris-HCl pH 7.5, 2 mM DTT, 20 µg/ml trypsin) followed by a second elution for 5 min with elution buffer 2 (2 M Urea, 50 mM Tris-HCl pH 7.5, 10 mM Chloroacetamide). Both eluates were combined and further incubated at room temperature over-night. Tryptic peptide mixtures were acidified to 1% TFA and loaded on Evotips (Evosep, Odense, Denmark).

### LC-MS analysis of on bead digested samples

Peptides were separated on 15 cm, 150 μM ID columns packed with C18 beads (1.9 μm) (Pepsep) on an Evosep ONE HPLC applying the ‘30 samples per day’ method and injected via a CaptiveSpray source with a 10 μm emitter into a timsTOF pro mass spectrometer (Bruker) operated in PASEF mode (Meier et al., 2018). Briefly, the DDA-PASEF scan range for both MS and MS/MS was set to 100 - 1700 m/z, and TIMS mobility range to 0.6 – 1.6 (V cm^−2^). TIMS ramp and accumulation times were set to 100 ms each, and 10 PASEF ramps recorded for a total cycle time of 1.17sec. MS/MS target intensity and intensity threshold were set to 20.000 and 1.000, respectively. An exclusion list of 0.4 min for precursors within 0.015 m/z and 0.015 V cm^−2^ width was also activated.

### Proximity labeling proteomics data analysis

Raw mass spectrometry data were analyzed with MaxQuant (v1.6.15.0) (Tyanova et al., 2016). Peak lists were searched against a manually curated version of the *Petunia inflata* FASTA database from Solgenomics.com, combined with 262 common contaminants by the integrated Andromeda search engine. False discovery rate was set to 1 % for both peptides (minimum length of 7 amino acids) and proteins. “Match between runs” (MBR) was enabled with a Match time window of 0.7, and a Match ion mobility window of 0.05 min. The proteinGroups.txt file was loaded in R and the data matrix filtered to remove proteins “only identified by site”, “reverse” and “potential contaminants”. The filtered matrix was analysed with R package DEP (Zhang et al., 2018). First, proteins were filtered to keep only proteins quantified in all three replicates in at least one (high stringency filtering) and subsequently missing values were imputed using MinProb. Differential enrichment of protein was comparing bait and controls at different timepoints, with manual adjustment of p-values using Benjamini-Hochberg (Benjamini & Hochberg, 1995). Significantly enriched proteins were obtained by an FDR of 0.05 and log2 fold change of 1.5. Volcano plots, PCA plots and iBAQ protein abundance ranks were plotted in RStudio (version 2023.06.0+421) and R (version 4.2.2), using proteomics datasets generated by DEP.

## AKNOWLEDGMENTS

We thank Ronald Koes and Francesca Quattrocchio from UvA, for providing protocols, *Petunia inflata* seeds, plasmid material and useful discussions. Mass spectrometry analyses were performed by the Proteomics Research Infrastructure (PRI) at the University of Copenhagen (UCPH), supported by the Novo Nordisk Foundation (NNF) (grant agreement number NNF19SA0059305). This research was funded by an Emerging Investigator Grant from NNF to TL (grant agreement number NNF19OC0055356).

