## Supplementary material for "Global organization of phenylpropanoid and anthocyanin pathways revealed by proximity labeling of trans-cinnamic acid 4-hydroxylase (CYP73A412) in *Petunia inflata* petal protoplasts"

**B**

**A**

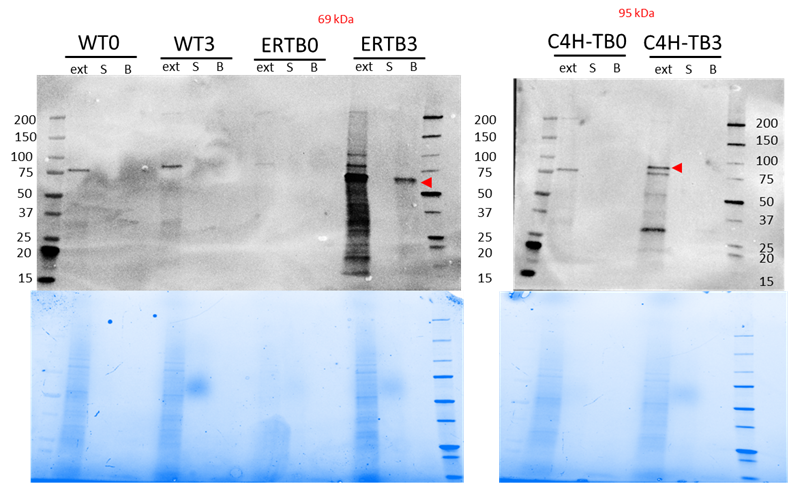

**Supplementary Figure 1**. Immunoblots to monitor affinity purification step. Protoplasts untransformed (WT) and transformed with plasmids for expression of ER-TurboID-EGFP (ERTB), CYP73A412-TurboID (C4H-TB) and untransformed (WT) were incubated with 50 µM biotin for 0 and 3 hours. After several washing steps, protoplasts were lysed and total protein was isolated. Biotinylated proteins were affinity purified using streptavidin coated beads. **A)** Immunoblot of the control samples WT and ERTB at 0 and 3 hours after incubation. **B)** Immunoblot for samples expressing CYP73-TurboID (C4H-TB). Lanes: ext: total protein extract, S: supernatant collected after affinity purification. B: proteins eluted from the beads after affinity purification. Red arrows indicate the band corresponding to “self” biotinylation.

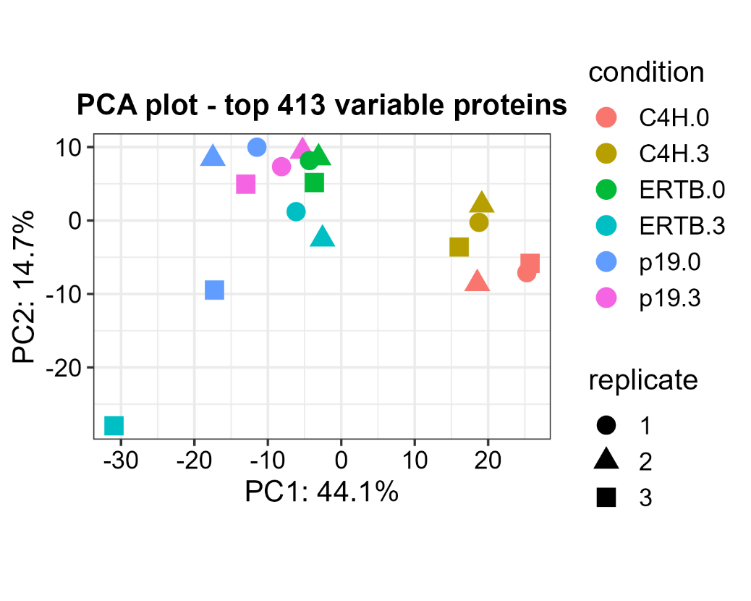

**Supplementary Figure 2.** Principal component analysis of proximity labeling experiments. C4H = CYP73

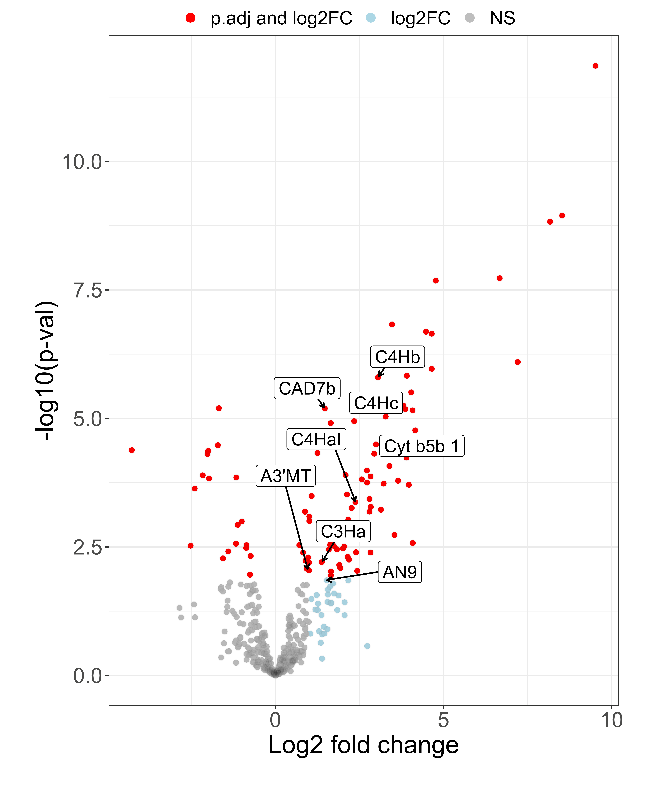

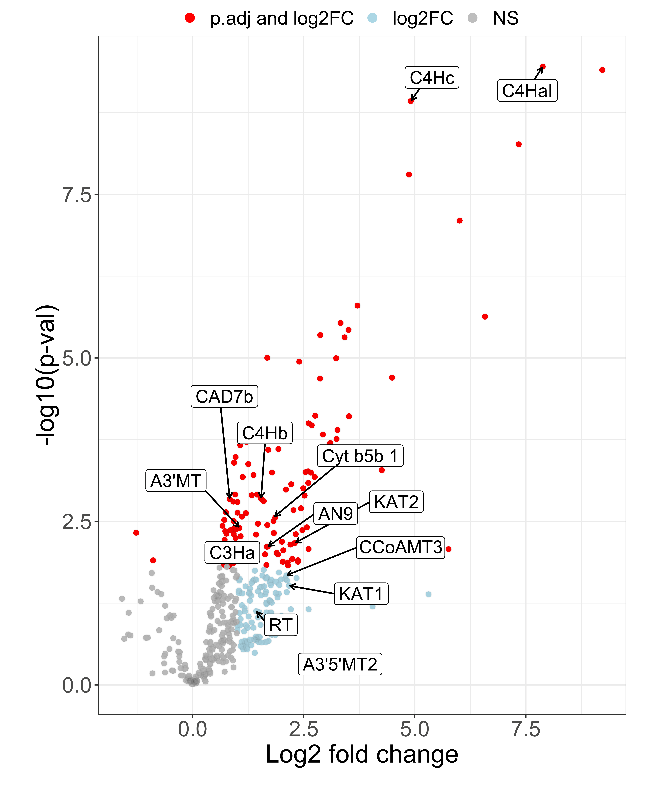

**Supplementary Figure 3.** Volcano plots of differentially enriched proteins in **A)** CYP73A412-TurboID vs ER-TurboID-EGFP at t=0 and **B)** CYP73A412-TurboID vs ER-TurboID-EGFP at t=180 min. Abbreviated enzyme names are indicated with boxes: A3’MT: Anthocyanin 3’-methyltransferase (MT/A3'MT); A3’5’MT2: Anthocyanin 3’5’-methyltransferase (MF2/A3'5’MT2); AN9: glutathione transferase; C4HaI, C4Hb, C4Hc: CYP73A paralogs; C3Ha: cinnamate 3-hydroxylase a; CAD7: cinnamyl alcohol dehydrogenase 7a; CCoAMT3: Caffeoyl-CoA O-methyltransferase 3; Cyt b5b1: Cytochrome b5b 1; KAT1: 3-ketoacyl-CoA thiolase 1; KAT2: 3-ketoacyl-CoA thiolase 2; RT: anthocyanin rhamnosyltransferase.

**Table1.** List of primers used for generation of vectors for proximity labeling

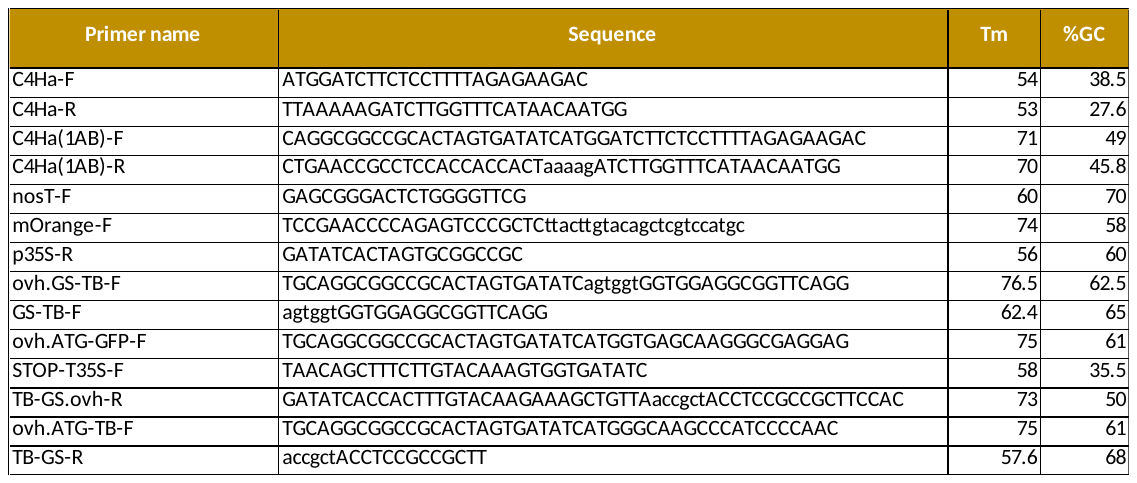

**Table 2.** List of significantly enriched proteins in CYP73A412-TurboID vs WT samples at timepoint t=0. Significance cutoff: adjusted p-value <0.05.

| **ID** | **Annotated protein name** | **log2 fold-change** |
| --- | --- | --- |
| Bait | CYP73A412-TurboID | 10.8 |
| Peinf101Scf00763g05002.1 | Sterol 14-demethylase (CYP51) | 8.78 |
| Peinf101Scf01631g01033.1 | Nodulin-related protein 1 | 7.87 |
| Peinf101Scf02326g01044.1 | Protein PAT1 homolog 2 | 6.87 |
| Peinf101Scf00838g06013.1 | NA | 6.59 |
| Peinf101Scf00151g10003.1 | Heme-binding-like protein At3g10130, chloroplastic | 6.23 |
| Peinf101Scf01857g02017.1 | Hsp70-Hsp90 organizing protein | 6.22 |
| Peinf101Scf01166g07038.1 | Phospholipase D beta 1 | 5.49 |
| Peinf101Scf00251g13014.1 | Fructose-bisphosphate aldolase 6, cytosolic | 4.83 |
| Peinf101Scf00951g08008.1 | Trans-cinnamate 4-monooxygenase C4H1 | 4.68 |
| Peinf101Scf02980g00010.1 | Cycloartenol-C-24-methyltransferase 1 | 4.63 |
| Peinf101Scf00175g15013.1 | Calcium-dependent lipid-binding protein | 4.59 |
| Peinf101Scf00554g06001.1 | Pyruvate kinase, cytosolic isozyme | 4.52 |
| Peinf101Scf00594g17025.1 | VAMP-like protein YKT61 | 4.29 |
| Peinf101Scf00633g00018.1 | Probable linoleate 9S-lipoxygenase 5 | 4.27 |
| Peinf101Scf00022g04043.1 | Protein HOMOLOG OF MAMMALIAN LYST-INTERACTING PROTEIN 5 | 4.26 |
| Peinf101Scf01077g02016.1 | Probable protein phosphatase 2C 58 | 4.14 |
| Peinf101Scf01163g00015.1 | Calcium-transporting ATPase 2, plasma membrane-type | 4.12 |
| Peinf101Scf03806g00033.1 | Trans-cinnamate 4-monooxygenase C4H2 | 4.09 |
| Peinf101Scf01786g07035.1 | Protein ROOT HAIR DEFECTIVE 3 | 4 |
| Peinf101Scf01482g05031.1 | TOM1-like protein 1 | 3.98 |
| Peinf101Scf03608g00001.1 | 3-ketoacyl-CoA synthase 3 | 3.91 |
| Peinf101Scf01047g05009.1 | NA | 3.83 |
| Peinf101Scf00686g01014.1 | Pleiotropic drug resistance protein 1 | 3.82 |
| Peinf101Scf00635g00007.1 | Cytochrome P450 71A6 | 3.81 |
| Peinf101Scf00091g29001.1 | Prohibitin-3, mitochondrial | 3.8 |
| Peinf101Scf00264g06017.1 | Peroxisome biogenesis protein 22 | 3.79 |
| Peinf101Scf01057g02013.1 | NA | 3.73 |
| Peinf101Scf01189g04042.1 | (+)-neomenthol dehydrogenase | 3.72 |
| Peinf101Scf00199g01006.1 | Cytochrome P450 76A2 | 3.71 |
| Peinf101Scf01047g04003.1 | Adhesive plaque matrix protein | 3.62 |
| Peinf101Scf01468g05003.1 | 24-methylenesterol C-methyltransferase 2 | 3.62 |
| Peinf101Scf00516g06012.1 | Cytochrome b5 | 3.57 |
| Peinf101Scf00145g20012.1 | Elongation factor 1-beta 1 | 3.56 |
| Peinf101Scf00471g10021.1 | Enoyl-CoA hydratase 2, peroxisomal | 3.53 |
| Peinf101Scf00261g12024.1 | Acetyl-coenzyme A carboxylase carboxyl transferase subunit beta, chloroplastic | 3.48 |
| Peinf101Scf02326g00003.1 | Reticulon-like protein B8 | 3.47 |
| Peinf101Scf06610g00047.1 | ER membrane protein complex subunit 4 | 3.45 |
| Peinf101Scf00655g08010.1 | ABC transporter E family member 2 | 3.31 |
| Peinf101Scf00670g05005.1 | Wax ester synthase/diacylglycerol acyltransferase 11 | 3.31 |
| Peinf101Ctg13285958g00001.1 | Cytosolic sulfotransferase 5 | 3.28 |
| Peinf101Scf00300g11006.1 | Heat shock cognate 70 kDa protein | 3.17 |
| Peinf101Scf00276g12026.1 | 60S ribosomal protein L13-1 | 3.16 |
| Peinf101Scf02582g00020.1 | Pyrophosphate-energized vacuolar membrane proton pump | 3.16 |
| Peinf101Scf00500g10031.1 | Hypersensitive-induced reaction 1 protein | 3.1 |
| Peinf101Scf00495g06008.1 | Pyrophosphate--fructose 6-phosphate 1-phosphotransferase subunit beta | 3.06 |
| Peinf101Scf00650g27060.1 | Dynamin-related protein 3A | 3.06 |
| Peinf101Scf01317g04052.1 | Late embryogenesis abundant protein Lea14-A | 3.02 |
| Peinf101Scf00650g23044.1 | Actin-11 | 3.01 |
| Peinf101Scf00653g11005.1 | V-type proton ATPase subunit d2 | 2.97 |
| Peinf101Scf00345g11024.1 | Mitochondrial outer membrane protein porin 2 | 2.91 |
| Peinf101Scf00659g02022.1 | Mitogen-activated protein kinase 9 | 2.89 |
| Peinf101Scf07097g00008.1 | GTP-binding protein YPTM2 | 2.89 |
| Peinf101Scf00559g15011.1 | Plasma membrane-associated cation-binding protein 1 | 2.87 |
| Peinf101Scf19669g00008.1 | Photosystem II D2 protein | 2.85 |
| Peinf101Scf00686g01027.1 | Pleiotropic drug resistance protein 1 | 2.83 |
| Peinf101Scf04640g00025.1 | Pleiotropic drug resistance protein 1 | 2.82 |
| Peinf101Scf02279g01029.1 | Non-symbiotic hemoglobin 1 | 2.8 |
| Peinf101Scf00179g07010.1 | Glucose-6-phosphate isomerase, cytosolic | 2.72 |
| Peinf101Scf00665g05023.1 | Uridine 5'-monophosphate synthase | 2.7 |
| Peinf101Scf00231g17017.1 | Mitochondrial outer membrane protein porin of 36 kDa | 2.63 |
| Peinf101Scf01267g04029.1 | Prohibitin-1, mitochondrial | 2.62 |
| Peinf101Scf03915g00017.1 | 25.3 kDa vesicle transport protein | 2.62 |
| Peinf101Scf00655g11011.1 | AP-4 complex subunit epsilon | 2.62 |
| Peinf101Scf00168g01001.1 | Prohibitin-1, mitochondrial | 2.61 |
| Peinf101Scf00058g12012.1 | Guanylate-binding protein 4 | 2.6 |
| Peinf101Scf00260g12014.1 | 5-methyltetrahydropteroyltriglutamate--homocysteine methyltransferase | 2.6 |
| Peinf101Scf01109g04006.1 | Ras-related protein RABB1c | 2.56 |
| Peinf101Scf00445g23011.1 | SEC12-like protein 1 | 2.54 |
| Peinf101Scf01044g06073.1 | Geraniol 8-hydroxylase | 2.54 |
| Peinf101Scf01871g00007.1 | Extradiol ring-cleavage dioxygenase | 2.53 |
| Peinf101Scf01468g08049.1 | SEC1 family transport protein SLY1 | 2.5 |
| Peinf101Scf00373g35002.1 | Mitochondrial import receptor subunit TOM40-1 | 2.49 |
| Peinf101Scf07318g00012.1 | ABC transporter C family member 2 | 2.47 |
| Peinf101Scf00120g02022.1 | Branched-chain-amino-acid aminotransferase-like protein 2 | 2.45 |
| Peinf101Scf00914g07043.1 | Ras-related protein RABA2a | 2.44 |
| Peinf101Scf02714g00035.1 | 60S ribosomal protein L3 | 2.39 |
| Peinf101Scf01294g01013.1 | 60S ribosomal protein L21-2 | 2.38 |
| Peinf101Scf01077g00021.1 | 26S proteasome non-ATPase regulatory subunit 2 homolog B | 2.37 |
| Peinf101Scf00889g13044.1 | T-complex protein 1 subunit epsilon | 2.36 |
| Peinf101Scf00434g11009.1 | Cytochrome P450 89A9 | 2.34 |
| Peinf101Scf03029g01029.1 | Ubiquitin carboxyl-terminal hydrolase 6 | 2.32 |
| Peinf101Scf00091g22003.1 | Ras-related protein Rab11D | 2.29 |
| Peinf101Scf00995g01001.1 | Nodulin-related protein 1 | 2.29 |
| Peinf101Scf00199g15006.1 | Enolase | 2.28 |
| Peinf101Scf00506g14021.1 | Cytochrome P450 98A2 | 2.27 |
| Peinf101Scf01380g00034.1 | Plasma membrane ATPase 3 | 2.25 |
| Peinf101Scf00232g01016.1 | Dolichyl-diphosphooligosaccharide--protein glycosyltransferase subunit 2 | 2.24 |
| Peinf101Scf00889g03034.1 | Glucose-6-phosphate 1-dehydrogenase, chloroplastic | 2.23 |
| Peinf101Scf00229g16011.1 | Poly(U)-specific endoribonuclease-B | 2.22 |
| Peinf101Scf01889g16040.1 | Alanine aminotransferase 2, mitochondrial | 2.21 |
| Peinf101Scf01786g09049.1 | 60S ribosomal protein L28-1 | 2.2 |
| Peinf101Scf01271g05017.1 | Caffeoyl-CoA O-methyltransferase 3 | 2.17 |
| Peinf101Scf00009g29006.1 | Coatomer subunit delta | 2.16 |
| Peinf101Ctg12523629g00001.1 | Cytosolic sulfotransferase 3 | 2.09 |
| Peinf101Scf02395g03053.1 | Inositol hexakisphosphate and diphosphoinositol-pentakisphosphate kinase VIP2 | 2.06 |
| Peinf101Scf06119g00002.1 | L-ascorbate peroxidase 2, cytosolic | 2.02 |
| Peinf101Scf00007g06012.1 | 14-3-3-like protein C | 1.96 |
| Peinf101Scf00303g09020.1 | Nuclear transport factor 2B | 1.96 |
| Peinf101Scf00337g05037.1 | Mitochondrial outer membrane protein porin 2 | 1.96 |
| Peinf101Scf01997g03024.1 | C-terminal binding protein AN | 1.96 |
| Peinf101Scf00364g12021.1 | Transketolase, chloroplastic | 1.94 |
| Peinf101Scf01996g00019.1 | Biotin carboxylase 1, chloroplastic | 1.94 |
| Peinf101Scf00043g02003.1 | Geraniol 8-hydroxylase | 1.94 |
| Peinf101Scf02312g02025.1 | ABC transporter G family member 36 | 1.91 |
| Peinf101Scf02344g00007.1 | Formate--tetrahydrofolate ligase | 1.9 |
| Peinf101Scf00170g03001.1 | UDP-glucose 6-dehydrogenase 1 | 1.89 |
| Peinf101Scf06265g00022.1 | 60S ribosomal protein L14-1 | 1.88 |
| Peinf101Scf01050g01004.1 | Very-long-chain enoyl-CoA reductase | 1.86 |
| Peinf101Scf04432g00003.1 | Lysine--tRNA ligase | 1.84 |
| Peinf101Scf00113g06019.1 | Ras-related protein RABE1c | 1.83 |
| Peinf101Scf00861g02014.1 | Glutathione S-transferase F11 | 1.8 |
| Peinf101Scf00947g08014.1 | Glutamine synthetase cytosolic isozyme 1-1 | 1.8 |
| Peinf101Scf00969g02020.1 | Transaldolase | 1.78 |
| Peinf101Scf00306g02009.1 | Pre-mRNA-processing protein 40A | 1.77 |
| Peinf101Scf02279g04046.1 | D-3-phosphoglycerate dehydrogenase 1, chloroplastic | 1.71 |
| Peinf101Scf00296g05002.1 | Cytochrome P450 CYP73A41200 | 1.7 |
| Peinf101Scf00665g07032.1 | Trifunctional UDP-glucose 4,6-dehydratase/UDP-4-keto-6-deoxy-D-glucose 3,5-epimerase/UDP-4-keto-L-rhamnose-reductase RHM1 | 1.7 |
| Peinf101Scf01662g01008.1 | 8-hydroxygeraniol dehydrogenase | 1.63 |
| Peinf101Scf01889g10037.1 | Heat shock 70 kDa protein 15 | 1.63 |
| Peinf101Scf03526g03027.1 | Regulator of nonsense transcripts 1 homolog | 1.63 |
| Peinf101Scf12523g00004.1 | Pyruvate kinase, cytosolic isozyme | 1.63 |
| Peinf101Scf04103g00072.1 | Nucleosome assembly protein 1;4 | 1.62 |
| Peinf101Scf01533g01038.1 | Alcohol dehydrogenase-like 3 | 1.59 |
| Peinf101Scf00763g01022.1 | ADP-ribosylation factor 2 | 1.57 |
| Peinf101Scf00974g19002.1 | Mitochondrial outer membrane protein porin of 34 kDa | 1.56 |
| Peinf101Scf01848g00038.1 | NA | 1.54 |
| Peinf101Scf00791g07004.1 | Phospho-2-dehydro-3-deoxyheptonate aldolase 2, chloroplastic | 1.52 |
| Peinf101Scf00060g09020.1 | Aspartate aminotransferase 3, chloroplastic | 1.48 |
| Peinf101Scf02318g00010.1 | Peptidyl-prolyl cis-trans isomerase | 1.48 |
| Peinf101Scf00400g05023.1 | Flavonoid 3',5'-methyltransferase | 1.42 |
| Peinf101Scf02093g03016.1 | Flavonoid 3',5'-methyltransferase | 1.41 |
| Peinf101Scf01267g01040.1 | Ketol-acid reductoisomerase, chloroplastic | 1.41 |
| Peinf101Scf01057g03044.1 | L-arabinokinase | 1.4 |
| Peinf101Scf00232g08024.1 | Alpha-aminoadipic semialdehyde synthase | 1.35 |
| Peinf101Scf00487g04019.1 | Pyruvate kinase 1, cytosolic | 1.31 |
| Peinf101Scf02085g05006.1 | Peroxisomal acyl-coenzyme A oxidase 1 | 1.29 |
| Peinf101Scf03119g00020.1 | Uridine kinase-like protein 1, chloroplastic | 1.25 |
| Peinf101Scf02092g00024.1 | Dynamin-related protein 5A | 1.24 |
| Peinf101Scf00232g10046.1 | Phosphoglycerate kinase, chloroplastic | 1.24 |
| Peinf101Scf00073g04027.1 | Dihydroflavonol 4-reductase | 1.16 |
| Peinf101Scf00261g14009.1 | 60S ribosomal protein L23a | 1.16 |
| Peinf101Scf01804g00023.1 | Phragmoplastin DRP1E | 1.15 |
| Peinf101Scf00936g05013.1 | Catalase isozyme 3 | 1.14 |
| Peinf101Scf00774g05003.1 | 2-methylpropanoate--CoA ligase CCL4 | 1.12 |
| Peinf101Scf01177g00002.1 | Secoisolariciresinol dehydrogenase | 1.05 |
| Peinf101Scf00364g03021.1 | Serine hydroxymethyltransferase 4 | 0.991 |
| Peinf101Scf00076g14015.1 | Phragmoplastin DRP1E | 0.981 |
| Peinf101Scf00471g00013.1 | Eukaryotic initiation factor 4A-11 | 0.93 |
| Peinf101Scf01192g02040.1 | Mitochondrial phosphate carrier protein 3, mitochondrial | 0.926 |
| Peinf101Scf00071g09027.1 | Heat shock protein 90-2 | 0.883 |
| Peinf101Scf03242g00011.1 | ADP,ATP carrier protein, mitochondrial | 0.858 |
| Peinf101Scf00665g25043.1 | Dynamin-2A | 0.781 |
| Peinf101Scf00665g22035.1 | Elongation factor 1-gamma 2 | 0.754 |
| Peinf101Scf03230g00022.1 | ADP,ATP carrier protein, mitochondrial | 0.72 |
| Peinf101Scf00055g14022.1 | Proteasome activator subunit 4 | 0.659 |
| Peinf101Scf00049g07035.1 | 60S ribosomal protein L3 | 0.57 |
| Peinf101Scf00256g03013.1 | Coatomer subunit beta'-2 | -0.663 |
| Peinf101Scf00364g03010.1 | Adenosylhomocysteinase | -0.708 |
| Peinf101Scf00082g01011.1 | 40S ribosomal protein SA | -0.721 |
| Peinf101Scf02446g00020.1 | Bifunctional L-3-cyanoalanine synthase/cysteine synthase 1, mitochondrial | -0.952 |
| Peinf101Scf00030g05012.1 | Glycine dehydrogenase (decarboxylating), mitochondrial | -0.989 |
| Peinf101Scf00633g07043.1 | 60S ribosomal protein L18-2 | -1.06 |
| Peinf101Scf00264g09029.1 | Aconitate hydratase, cytoplasmic | -1.11 |
| Peinf101Scf00423g05008.1 | 60S ribosomal protein L32-1 | -1.12 |
| Peinf101Scf01367g05017.1 | Probable nucleolar protein 5-2 | -1.19 |
| Peinf101Scf00809g06022.1 | Eukaryotic translation initiation factor 5B | -1.23 |
| Peinf101Scf01427g04015.1 | Succinate dehydrogenase [ubiquinone] flavoprotein subunit 1, mitochondrial | -1.29 |
| Peinf101Scf01734g02016.1 | 60S ribosomal protein L13a-4 | -1.32 |
| Peinf101Scf02187g02012.1 | Sister chromatid cohesion protein PDS5 homolog B-B | -1.32 |
| Peinf101Scf10642g00004.1 | Proteasome subunit beta type-4 | -1.45 |
| Peinf101Scf02381g00005.1 | Succinate--CoA ligase [ADP-forming] subunit beta, mitochondrial | -1.61 |
| Peinf101Scf00251g19015.1 | Monodehydroascorbate reductase | -1.84 |
| Peinf101Scf02485g01008.1 | Histone H4 | -1.9 |
| Peinf101Ctg13480496g00003.1 | Nucleolin 2 | -1.95 |
| Peinf101Scf00199g16014.1 | Histone deacetylase HDT1 | -1.96 |
| Peinf101Scf00435g05001.1 | H/ACA ribonucleoprotein complex subunit 4 | -1.97 |
| Peinf101Scf00766g02012.1 | Isocitrate dehydrogenase [NADP] | -1.98 |
| Peinf101Scf07361g00027.1 | Formate dehydrogenase, mitochondrial | -2.06 |
| Peinf101Scf01186g06029.1 | Nucleolar protein 56 | -2.1 |
| Peinf101Scf00442g02021.1 | Aspartate aminotransferase, mitochondrial | -2.31 |
| Peinf101Scf00672g10011.1 | Malate dehydrogenase | -2.69 |
| Peinf101Scf00351g04016.1 | Aminomethyltransferase, mitochondrial | -2.81 |
| Peinf101Scf01878g01003.1 | Histone H3.2 | -3.61 |
| Peinf101Scf00650g13039.1 | Histone H2A.1 | -4.07 |

**Table 3.** List of significantly enriched proteins in CYP73A412-TurboID vs WT at timepoint = 3h. Significance cutoff: adjusted p-value < 0.05

| **ID** | **Annotated protein name** | **log2 fold-change** |
| --- | --- | --- |
| Peinf101Scf01142g01025.1 | Protein FAR1-RELATED SEQUENCE 6 | 8.63 |
| Peinf101Scf01857g02017.1 | Hsp70-Hsp90 organizing protein | 7.48 |
| Peinf101Scf00951g08008.1 | Trans-cinnamate 4-monooxygenase C4H1 | 7.08 |
| Peinf101Scf00838g06013.1 | NA | 7.05 |
| Peinf101Scf01631g01033.1 | Nodulin-related protein 1 | 6.81 |
| Peinf101Scf00022g04043.1 | Protein HOMOLOG OF MAMMALIAN LYST-INTERACTING PROTEIN 5 | 4.96 |
| Peinf101Scf00296g05002.1 | Cytochrome P450 CYP73A41200 | 4.91 |
| Peinf101Scf01166g07038.1 | Phospholipase D beta 1 | 4.56 |
| Peinf101Scf00554g06001.1 | Pyruvate kinase, cytosolic isozyme | 4.43 |
| Bait | CYP73A412-TurboID | 4.03 |
| Peinf101Scf00655g08010.1 | ABC transporter E family member 2 | 3.72 |
| Peinf101Scf00633g00018.1 | Probable linoleate 9S-lipoxygenase 5 | 3.71 |
| Peinf101Scf00264g06017.1 | Peroxisome biogenesis protein 22 | 3.65 |
| Peinf101Scf00303g09020.1 | Nuclear transport factor 2B | 3.48 |
| Peinf101Scf01077g02016.1 | Probable protein phosphatase 2C 58 | 3.48 |
| Peinf101Scf00345g11024.1 | Mitochondrial outer membrane protein porin 2 | 3.42 |
| Peinf101Scf00231g17017.1 | Mitochondrial outer membrane protein porin of 36 kDa | 3.39 |
| Peinf101Scf01889g16040.1 | Alanine aminotransferase 2, mitochondrial | 3.36 |
| Peinf101Scf00229g16011.1 | Poly(U)-specific endoribonuclease-B | 3.25 |
| Peinf101Scf00175g15013.1 | Calcium-dependent lipid-binding protein | 3.16 |
| Peinf101Scf01047g04003.1 | Adhesive plaque matrix protein | 3.14 |
| Peinf101Scf01633g10042.1 | Respiratory burst oxidase homolog protein C | 3.04 |
| Peinf101Scf01294g01013.1 | 60S ribosomal protein L21-2 | 3.04 |
| Peinf101Scf02326g00003.1 | Reticulon-like protein B8 | 3.02 |
| Peinf101Ctg12523629g00001.1 | Cytosolic sulfotransferase 3 | 3 |
| Peinf101Scf00655g11011.1 | AP-4 complex subunit epsilon | 3 |
| Peinf101Scf00995g01001.1 | Nodulin-related protein 1 | 2.97 |
| Peinf101Scf01853g03035.1 | Nucleosome assembly protein 1;2 | 2.97 |
| Peinf101Scf01047g05009.1 | NA | 2.95 |
| Peinf101Scf00653g11005.1 | V-type proton ATPase subunit d2 | 2.91 |
| Peinf101Scf01271g05017.1 | Caffeoyl-CoA O-methyltransferase 3 | 2.8 |
| Peinf101Scf00559g15011.1 | Plasma membrane-associated cation-binding protein 1 | 2.66 |
| Peinf101Scf01482g05031.1 | TOM1-like protein 1 | 2.59 |
| Peinf101Scf00274g00004.1 | Chorismate synthase 1, chloroplastic | 2.57 |
| Peinf101Scf03029g01029.1 | Ubiquitin carboxyl-terminal hydrolase 6 | 2.56 |
| Peinf101Scf02395g03053.1 | Inositol hexakisphosphate and diphosphoinositol-pentakisphosphate kinase VIP2 | 2.54 |
| Peinf101Scf00300g11006.1 | Heat shock cognate 70 kDa protein | 2.39 |
| Peinf101Scf00665g05023.1 | Uridine 5'-monophosphate synthase | 2.39 |
| Peinf101Scf01317g04052.1 | Late embryogenesis abundant protein Lea14-A | 2.33 |
| Peinf101Scf01543g00042.1 | Trihelix transcription factor GT-3a | 2.28 |
| Peinf101Scf00120g02022.1 | Branched-chain-amino-acid aminotransferase-like protein 2 | 2.11 |
| Peinf101Scf02279g01029.1 | Non-symbiotic hemoglobin 1 | 2.07 |
| Peinf101Scf07318g00012.1 | ABC transporter C family member 2 | 2.06 |
| Peinf101Scf03526g03027.1 | Regulator of nonsense transcripts 1 homolog | 2.06 |
| Peinf101Scf00058g12012.1 | Guanylate-binding protein 4 | 2.02 |
| Peinf101Scf02980g00010.1 | Cycloartenol-C-24-methyltransferase 1 | 2.02 |
| Peinf101Scf00251g13014.1 | Fructose-bisphosphate aldolase 6, cytosolic | 1.98 |
| Peinf101Scf01786g07035.1 | Protein ROOT HAIR DEFECTIVE 3 | 1.98 |
| Peinf101Scf00306g02009.1 | Pre-mRNA-processing protein 40A | 1.97 |
| Peinf101Ctg13285958g00001.1 | Cytosolic sulfotransferase 5 | 1.94 |
| Peinf101Scf01163g00015.1 | Calcium-transporting ATPase 2, plasma membrane-type | 1.9 |
| Peinf101Scf03806g00033.1 | Trans-cinnamate 4-monooxygenase C4H2 | 1.84 |
| Peinf101Scf00516g06012.1 | Cytochrome b5 | 1.8 |
| Peinf101Scf00091g29001.1 | Prohibitin-3, mitochondrial | 1.72 |
| Peinf101Scf00686g01014.1 | Pleiotropic drug resistance protein 1 | 1.59 |
| Peinf101Scf01969g05037.1 | Uncharacterized protein At5g39570 | 1.47 |
| Peinf101Scf00506g14021.1 | Cytochrome P450 98A2 | 1.35 |
| Peinf101Scf00168g01001.1 | Prohibitin-1, mitochondrial | 1.03 |
| Peinf101Scf01533g01038.1 | Alcohol dehydrogenase-like 3 | 0.942 |
| Peinf101Scf01177g00002.1 | Secoisolariciresinol dehydrogenase | 0.746 |
| Peinf101Scf00061g05012.1 | Putative rRNA 2'-O-methyltransferase fibrillarin 3 | 0.666 |
| Peinf101Scf00633g07043.1 | 60S ribosomal protein L18-2 | -1.01 |
| Peinf101Scf02446g00020.1 | Bifunctional L-3-cyanoalanine synthase/cysteine synthase 1, mitochondrial | -1.01 |
| Peinf101Scf07361g00027.1 | Formate dehydrogenase, mitochondrial | -1.04 |
| Peinf101Scf00251g19015.1 | Monodehydroascorbate reductase | -1.08 |
| Peinf101Scf00672g10011.1 | Malate dehydrogenase | -1.57 |
| Peinf101Scf02381g00005.1 | Succinate--CoA ligase [ADP-forming] subunit beta, mitochondrial | -1.66 |
| Peinf101Scf10642g00004.1 | Proteasome subunit beta type-4 | -2.12 |
| Peinf101Scf00351g04016.1 | Aminomethyltransferase, mitochondrial | -2.88 |
